# Predicting viral host range for reverse zoonosis surveillance

**DOI:** 10.64898/2026.09.30.755793

**Authors:** Edward B. Irvine, Thomas Bikias, Théo Schifferli, Leon Plath, Jiami Han, NingNing Chen, Evangelos Stamkopoulos, Berk Sahin, Lester Frei, Joseph M. Taft, Sai T. Reddy

## Abstract

Zoonotic transmission of viruses has triggered major outbreaks throughout history. However, human-to-animal transmission, or “reverse zoonotic” transmission, also represents an underappreciated public health threat. Reverse zoonoses can establish novel animal reservoirs that fuel viral evolution and enable eventual spillback into humans, underscoring the importance of defining viral host range. Current approaches for host range prediction are limited. Experimental methods are retrospective and rely on brute-force testing in a small number of models, whereas computational approaches often lack experimental grounding. Here, we present a platform that integrates high-throughput experimental screening with deep learning to predict viral host range from sequence. We screened a yeast-displayed SARS-CoV-2 receptor-binding domain library against 15 angiotensin-converting enzyme 2 (ACE2) orthologs from diverse species, and deep sequenced the ACE2-binding and non-binding populations to generate an atlas of 8.8 million unique RBD:ACE2 interaction measurements. These data were used to train a transformer-based deep learning model that predicts RBD:ACE2 compatibility across animal species from sequence alone, and generalizes in a zero-shot manner to ACE2 orthologs not observed during training. Critically, the model pairs each prediction with a distance-based uncertainty estimate, allowing hundreds of candidate species to be triaged by both predicted host compatibility and confidence. This framework enables the rapid prioritization of animal species for SARS-CoV-2 surveillance and provides a general strategy for anticipating viral host range.

## INTRODUCTION

Zoonotic transfer of viruses from animals to humans is a primary cause of emerging infectious disease threats globally (Jones et al., 2008). This species-to-species transmission introduces viruses to new host populations, enabling evolutionary adaptation and rapid spread through immunologically naive communities. In recent history, severe viral epidemics driven by pathogens such as HIV, Ebola virus, and monkeypox virus, were each triggered by zoonotic events (Bunge et al., 2022; Gire et al., 2014; Hahn et al., 2000). Coronaviruses, in particular, have exhibited a remarkable proclivity to jump across species barriers, with bat-origin viruses believed to be the source of seasonal human coronaviruses (OC43, NL63, and 229E), as well as major epidemics including MERS-CoV, SARS-CoV-1, and SARS-CoV-2 (Boni et al., 2020; Corman et al., 2015, 2014; Huynh et al., 2012; Latinne et al., 2020; Lau et al., 2005; Li et al., 2005). Influenza H5N1 threatens to follow a similarly dangerous trajectory, with recent evidence of adaptation to mammalian hosts and occasional human infections (Capelastegui and Goldhill, 2025; Caserta et al., 2024).

While public and scientific attention understandably focuses on animal-to-human spillover, viral transmission is bidirectional, and human-to-animal spillover, or “reverse zoonoses”, represent a distinct and underappreciated evolutionary risk (Jia et al., 2021). Though traditionally a concern in veterinary medicine, reverse zoonoses are now recognized as potential drivers of viral adaptation with direct implications for human public health. For instance, numerous cases of SARS-CoV-2 transmission from humans to animals have now been documented across a range of species, including dogs, cats, ferrets, otters, gorillas, mink, and white-tailed deer (“SARS-CoV-2 in Animals,” n.d.), with subsequent intraspecies transmission reported in several species (Fagre et al., 2021; Freuling et al., 2020; Hale et al., 2022; Kim et al., 2020; Lu et al., 2021; Oude Munnink et al., 2021; Schlottau et al., 2020; Shi et al., 2020; Sia et al., 2020). Most notably, SARS-CoV-2 has spread extensively within the white-tailed deer and mink populations (Hale et al., 2022; Lu et al., 2021; Oude Munnink et al., 2021). In deer, animal-to-animal spread gave rise to mutations in the SARS-CoV-2 spike protein rarely observed in humans (Hale et al., 2022). Likewise, a spike protein mutation acquired in mink conferred increased viral escape capacity from both therapeutic monoclonal antibodies and convalescent plasma (Hoffmann et al., 2021). Notably, several mink farm workers were infected with strains bearing an animal sequence signature (Oude Munnink et al., 2021), demonstrating that reverse zoonosis can lead to the re-entry of animal-adapted viral variants into the human population.

These cases demonstrate that reverse zoonotic events can establish novel animal reservoirs for viral replication and evolution. While documented in these instances, such reservoirs may also develop silently in poorly surveyed animal species. Over time, these cryptic reservoirs may generate highly mutated variants that, upon re-entry into the human population, could circumvent existing immunity and trigger new waves of infection. This consistent threat of interspecies viral transmission highlights that a robust understanding of viral host range is fundamental to understanding viral evolution and predicting future outbreaks.

Current approaches to determine viral host range are limited. Experimental work relies on the brute-force testing of individual viral variants in a small number of *in vitro* and animal models (Conceicao et al., 2020; Li et al., 2024; Munster et al., 2020; Santini and Edwards, 2020; Shi et al., 2020; Sia et al., 2020), whereas computational approaches often lack experimental grounding (Babayan et al., 2018; Damas et al., 2020; Guo et al., 2021; Mock et al., 2021). *In silico* approaches also typically model host compatibility for a single, fixed viral variant, and cannot resolve how compatibility changes as the virus evolves – even though host range can be gained or lost through mutation (Dinnon et al., 2020; Halfmann et al., 2022; Starr et al., 2022; Thakur et al., 2022). Understanding these shifts requires models that capture variation on both sides of the interaction.

Here, we introduce a platform that integrates high-throughput experimental screening with deep learning to predict viral host range (**Fig. 1**). Specifically, we leveraged a yeast surface display library of the SARS-CoV-2 spike receptor-binding domain (RBD) (Frei et al., 2025), sorting it using a panel of 15 soluble angiotensin-converting enzyme 2 (ACE2) orthologs from diverse avian and mammalian species that span broad phylogenetic and sequence diversity. These sorted populations were deep sequenced to generate a high-resolution dataset that served as training data for a transformer-based deep learning model. The model accurately predicted ACE2 binding across the 15 species evaluated and generalized in a zero-shot manner to ACE2 orthologs not observed during training. By jointly embedding RBD and ACE2 sequences, the model can predict binding for any RBD variant against any animal with an available ACE2 sequence, while an integrated uncertainty metric flags predictions beyond the training distribution. This platform provides a tool for the rapid assessment of SARS-CoV-2 spillover risk simply using RBD and ACE2 sequence, offering an early warning system for animal reservoirs that could drive future waves of infection. This framework may be extended to other viral families and host receptors, offering a general approach to transform how we anticipate and respond to viral threats.

**Fig. 1.**
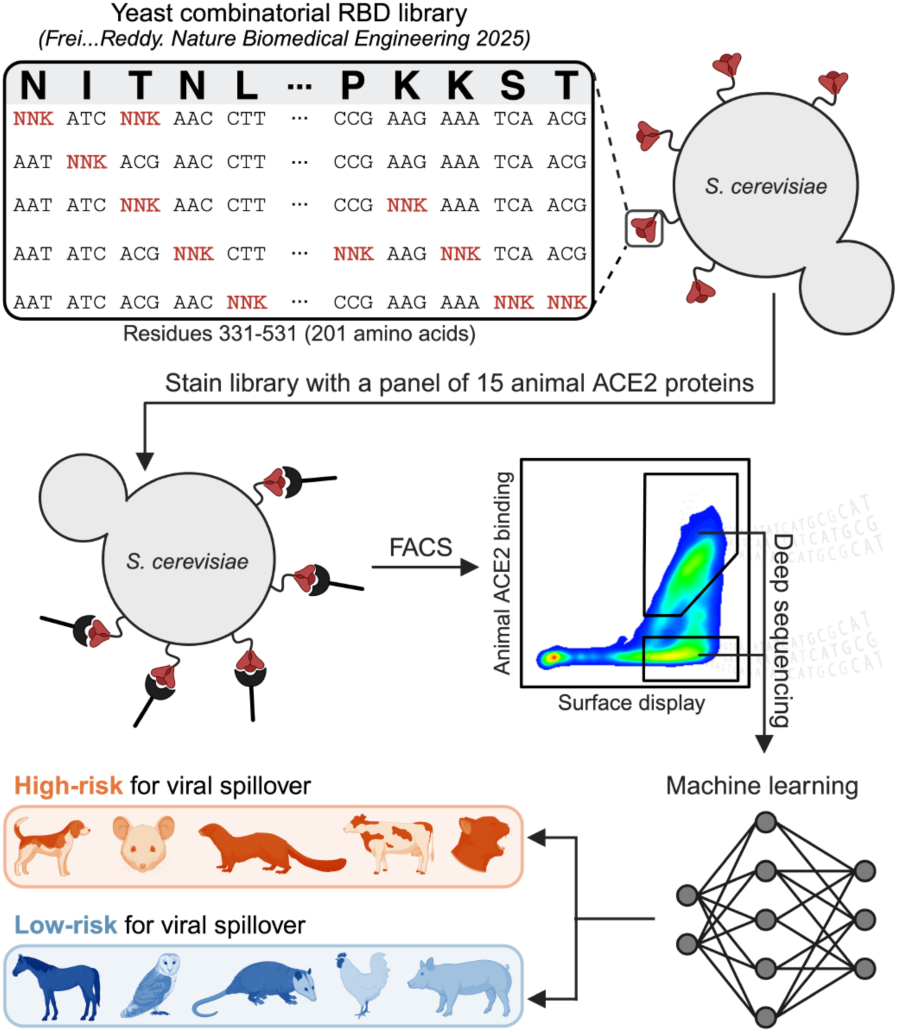
A machine learning-guided platform for predicting viral host range. A combinatorial yeast surface display library (diversity = 2×10^8^) covering residues 331–531 of the SARS-CoV-2 Omicron BA.1 spike RBD was stained with a panel of 15 ACE2 orthologs from diverse mammalian and avian species. Sorted populations were deep sequenced to generate a high-resolution dataset linking RBD variants to animal ACE2 binding. These data were used to train a deep learning model that predicts RBD:ACE2 interactions as a proxy for viral host range. Created with <u>BioRender.com</u>.

## RESULTS

### Rational selection of a diverse panel of ACE2 orthologs

A prerequisite for interspecies transmission is the ability of a virus to engage a compatible receptor in the new potential host. For SARS-CoV-2, host cell entry is mediated by the spike glycoprotein, which binds the angiotensin-converting enzyme 2 (ACE2) on the surface of target cells (V’kovski et al., 2021). As such, ACE2 compatibility is a primary determinant of SARS-CoV-2 host range. To establish a generalizable platform for SARS-CoV-2 host range prediction based on ACE2 binding, we first sought to assemble and characterize a panel of ACE2 proteins from a broad range of animal species.

We assembled a collection of 452 full-length ACE2 orthologs by combining protein sequences from NCBI and UniProt. This collection underwent manual curation to remove entries with poor annotation quality, incomplete coding regions, or substantial sequence truncations. The curated ACE2 sequence collection included representatives from all major vertebrate classes: mammals, birds, reptiles, amphibians, and fish (**Supplementary Table 1**). To visualize the diversity of these sequences across ACE2 sequence space, sequences were aligned using MAFFT (Katoh et al., 2002), one-hot encoded, and projected into low-dimensional space via principal component analysis (PCA) (**Fig. 2a**). ACE2 sequences clustered clearly by vertebrate class (**Fig. 2a**).

**Fig. 2.**
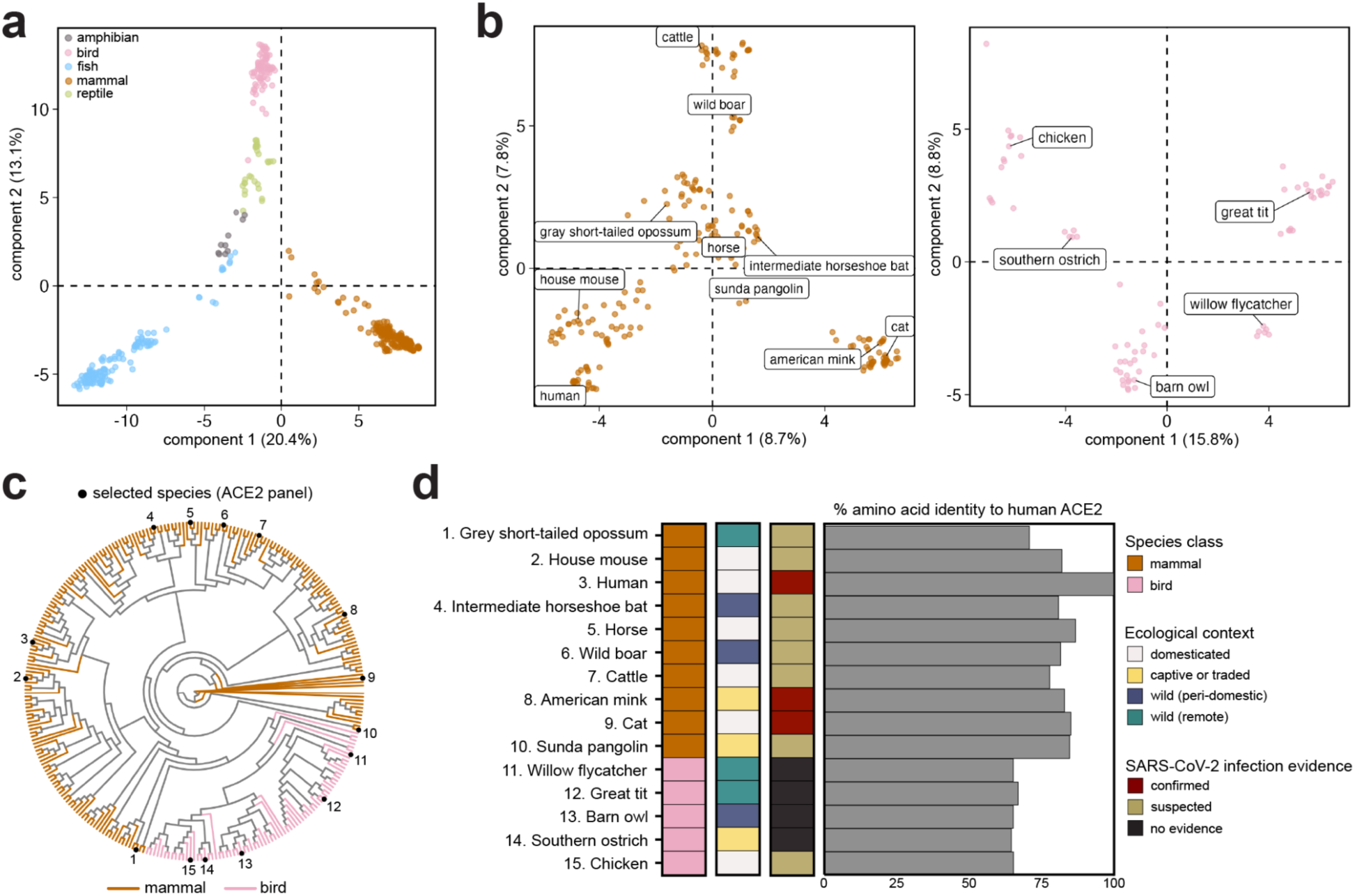
Rational selection of a diverse panel of animal ACE2 orthologs. **a,** Principal component analysis (PCA) of 452 animal ACE2 amino acid sequences, aligned using MAFFT and one-hot-encoded. **b,** PCA of ACE2 sequences subsetted by species class: mammals (left) and birds (right). The 15 ACE2 proteins selected for experimental analyses – ten from mammals and five from birds – are labeled. **c,** Maximum likelihood phylogenetic tree of mammal and bird ACE2 amino acid sequences. Tree branches are colored by species class. Black points indicate the 15 selected ACE2 orthologs. **d,** Metadata associated with the 15 selected ACE2 orthologs. From left to right: species class, ecological context, evidence of natural SARS-CoV-2 infection evidence (confirmed, suspected, or no evidence), and amino acid sequence identity to human ACE2. “Confirmed” indicates natural infection; “Suspected” refers to experimental infection or indirect evidence of susceptibility.

Given the reported susceptibility of mammals and birds to coronaviruses (V’kovski et al., 2021), we focused our downstream efforts on ACE2 orthologs from these two vertebrate classes. Sub-clustering within mammalian and avian ACE2 sequences revealed discrete groups likely reflecting evolutionary similarities (**Fig. 2b**). To maximize coverage of ACE2 sequence space within these classes, we selected 15 representative orthologs – ten from mammals and five from birds – that spanned the major clusters (**Fig. 2b**). To assess the phylogenetic breadth of the selected sequences, we constructed a maximum likelihood tree of the aligned mammalian and avian ACE2 sequences. The selected orthologs were broadly distributed across the tree, confirming the phylogenetic diversity of the panel (**Fig. 2c**).

In addition to sequence and phylogenetic diversity, the selected species capture a range of ecological contexts, from domesticated animals to remote wild species with minimal human contact (**Fig. 2d**). Among the 15 species, three have confirmed evidence of natural SARS-CoV-2 infection, eight have suspected or experimental evidence, and four have no reported evidence of infection or susceptibility (**Fig. 2d**) (Doliff and Martens, 2022; Lu et al., 2021; Oude Munnink et al., 2021). Interestingly, evidence of SARS-CoV-2 infection or susceptibility was not strictly ordered by amino acid identity to human ACE2 (**Fig. 2d**), suggesting that overall receptor similarity alone is insufficient to infer host compatibility. All ACE2 proteins were obtained commercially (either off-the-shelf or custom-produced), and SDS-PAGE confirmed expression and expected molecular weight for all custom-produced proteins (**Extended Data Fig. 1a,b**). Collectively, this work establishes a panel of sequence-diverse, phylogenetically broad, and ecologically varied ACE2 orthologs, providing a robust foundation for evaluating RBD:ACE2 compatibility across species.

### Construction of an atlas of RBD:ACE2 interactions across species

General-purpose protein-protein interaction models often lack the sensitivity required to accurately predict the effects of single (or few) amino acid substitutions on binding (Chakraborty et al., 2026; McMaster et al., 2026; Ullanat et al., 2026). Thus, to enable high-resolution predictive modeling of RBD:ACE2 interactions across species, we sought to generate a large-scale, domain-specific RBD:ACE2 interaction dataset experimentally.

We started with a previously published combinatorial mutagenesis library of the SARS-CoV-2 Omicron BA.1 RBD. The library spans the entire RBD (**Fig. 3a**), and contains an experimental diversity of 2.0×10^8^ variants with an average of 3.6 mutations per sequence. It was designed to comprehensively sample the mutational landscape of Omicron BA.1 while capturing higher-order epistatic interactions (Frei et al., 2025). We screened this library for binding across our panel of 15 diverse animal ACE2 proteins, isolating ACE2-binding and non-binding populations for each species through multiple rounds of fluorescence-activated cell sorting (FACS) (**Fig. 3b** and **Extended Data Fig. 2a**). The sorted populations were deep sequenced, pre-processed to remove low-quality reads and variants with fewer than two counts, and translated to amino acid sequences to generate clean datasets for each species. All animal species showed a higher proportion of non-binders than binders, with non-binders comprising 51.2 - 98.4% of sequenced variants across species, consistent with the pattern observed by flow cytometry (**Fig. 3c** and **Extended Data Fig. 2a**). Furthermore, many RBD variants had experimental binding data for only a single animal ACE2 protein (**Extended Data Fig. 2b**), as expected given that the throughput of FACS (∼10^7^ cells) undersamples the full library (∼10^8^ variants).

**Fig. 3.**
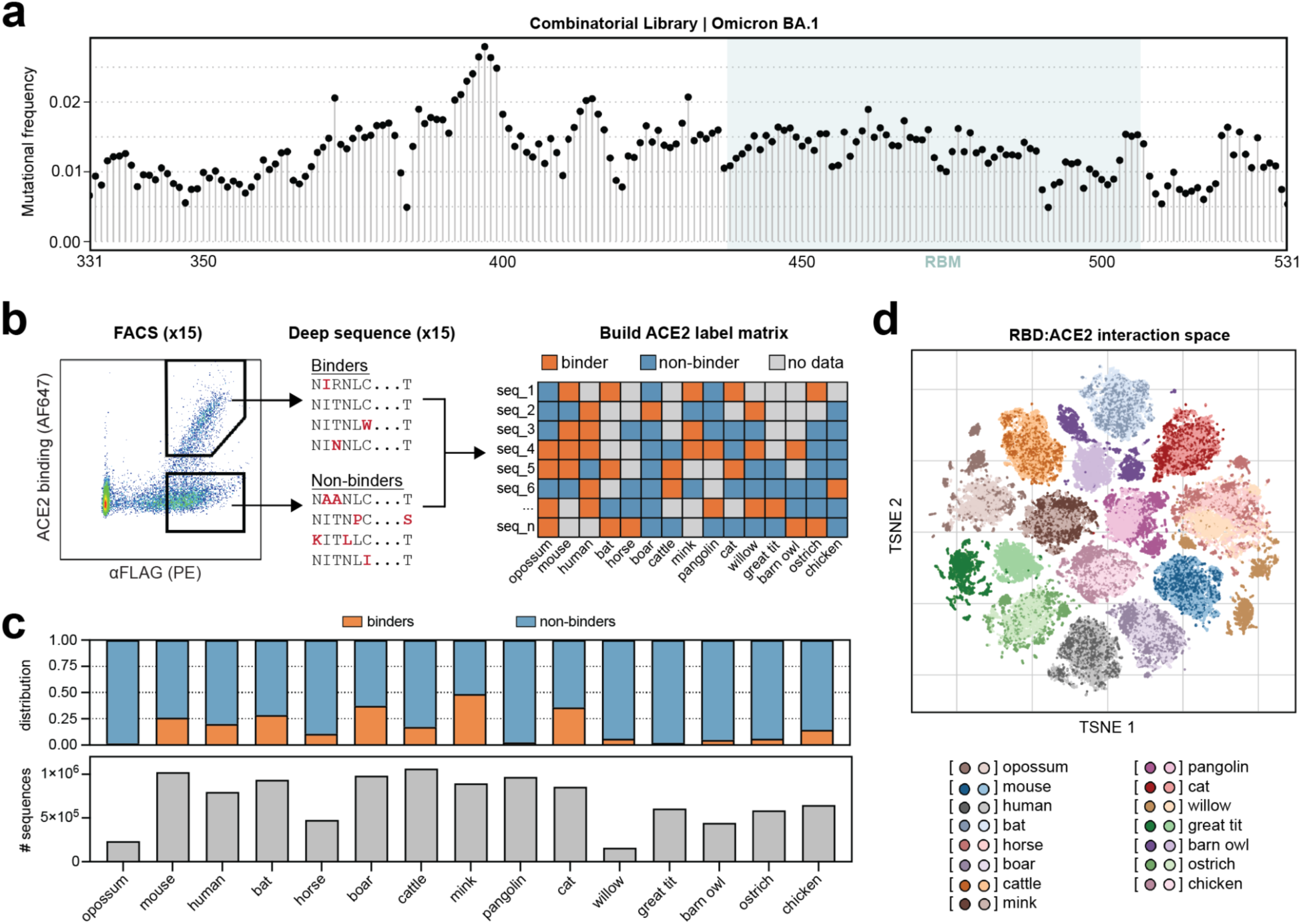
Construction of an atlas of RBD:ACE2 interactions across species. **a,** Frequency of mutations at each residue (331-531) in the Omicron BA.1 RBD combinatorial library. The receptor-binding motif (RBM) is highlighted in light blue. **b,** Binding and non-binding populations for each of the 15 animal ACE2 orthologs were isolated via FACS and deep sequenced, generating a training dataset where each RBD sequence is assigned as a binder (orange), non-binder (blue), or no data (grey) for each animal ACE2 ortholog. **c,** Number (bottom) and class distribution (top) of training data following pre-processing. Human, mouse, bat, and cattle ACE2 were screened at 50nM. All remaining orthologs were screened at 500nM to capture lower-affinity binders. Binder proportions are therefore not directly comparable across species. **d,** TSNE visualization of the RBD:ACE2 interaction space derived from concatenated ESM2 embeddings of RBD and ACE2 sequences, with points colored by species. For each species, darker points (left) indicate binders and lighter points (right) non-binders.

This effort resulted in a dataset of 8.8 million unique RBD:ACE2 interaction measurements spanning 15 animal species. To examine the composition of this dataset, we embedded RBD and ACE2 sequences using the ESM2 protein language model (Lin et al., 2023), concatenated their representations, and projected them into low-dimensional space (**Fig. 3d**). RBD:ACE2 sequence pairs robustly clustered by animal species. However, for most species, binding and non-binding sequence pairs did not clearly separate in embedding space (**Fig. 3d**), indicating that pretrained sequence representations alone do not resolve the determinants of RBD:ACE2 binding and highlighting the need for supervised modeling. Together, this work establishes a comprehensive atlas of RBD:ACE2 interactions across animal species, providing a large-scale training dataset for modeling viral receptor engagement across diverse hosts.

### Deep learning models accurately predict RBD:ACE2 binding

Previous work has shown that combining yeast display screening of SARS-CoV-2 RBD mutagenesis libraries with deep sequencing and machine learning – an approach known as deep mutational learning – enables the impact of combinatorial RBD mutations on antibody and human ACE2 binding to be accurately predicted (Frei et al., 2025; Shlesinger et al., 2025; Taft et al., 2022). Given the predictive power and clinical insight provided by these models, we reasoned that an analogous framework could be used to predict ACE2 engagement across diverse animal species, thus enabling prediction of cross-species RBD:ACE2 compatibility.

Using the RBD:ACE2 interaction dataset, we trained machine learning models to predict RBD:ACE2 binding across all 15 species directly from protein sequence. To systematically benchmark model performance, we evaluated a diverse set of modeling strategies, including gradient-boosted trees (XGBoost), a multilayer perceptron (MLP), a fine-tuned ESM-8M encoder, and a transformer-based architecture, providing a benchmark across distinct classes of sequence-based models (**Extended Data Fig. 3a**). Each model was trained using a cluster-based data split in which RBD variants were grouped by sequence similarity from one-hot encoded amino acid sequences into five distinct clusters using k-means clustering. Two of the five resulting clusters were used for training, while the remaining clusters were reserved for validation and testing to reduce sequence-level leakage and assess generalization to distinct mutational patterns. Model performance was assessed using Matthews correlation coefficient (MCC), which provides a robust measure of performance even in the presence of class imbalance (Chicco and Jurman, 2020). Among the approaches evaluated, the transformer achieved the highest average MCC across species and thus was selected as our primary model (**Extended Data Fig. 3a**).

Specifically, RBD and ACE2 sequences were embedded using the ESM2 protein language model (Lin et al., 2023), generating contextual amino acid representations for each sequence (**Fig. 4a**). To construct an interaction-aware representation for each RBD:ACE2 pair the RBD embeddings were retained at residue-level resolution, while the ACE2 embeddings were mean-pooled into a receptor-level representation to accommodate their greater and variable sequence lengths. The ACE2 representation was then broadcast across RBD positions and combined with each RBD residue embedding using three feature components: the original RBD residue embedding, the absolute difference between the RBD and ACE2 embeddings, and their element-wise product (**Fig. 4a**). Feature importance analysis supported the contribution of these interaction-derived terms to prediction, with the element-wise product providing the strongest non-RBD contribution (**Extended Data Fig. 3b**). The resulting features were projected into a shared latent space and processed with a transformer encoder, enabling the model to learn higher-order dependencies across the RBD sequence that are conditioned on ACE2 sequence (**Fig. 4a**). Encoded features were subsequently passed through a position-wise feedforward network and downstream classification head to predict RBD:ACE2 binding as a binary outcome (**Fig. 4a**). The model was trained using binary cross-entropy loss, with hyperparameters optimized on the validation set and early stopping used to limit overfitting. This architecture enables predictive modeling of RBD:ACE2 interactions across multiple species using a single unified model, rather than requiring separate models for each ACE2 ortholog. Ultimately, the model achieved robust performance across all 15 animals, with an average MCC of 0.830 on the held-out test set (**Fig. 4b**).

**Fig. 4.**
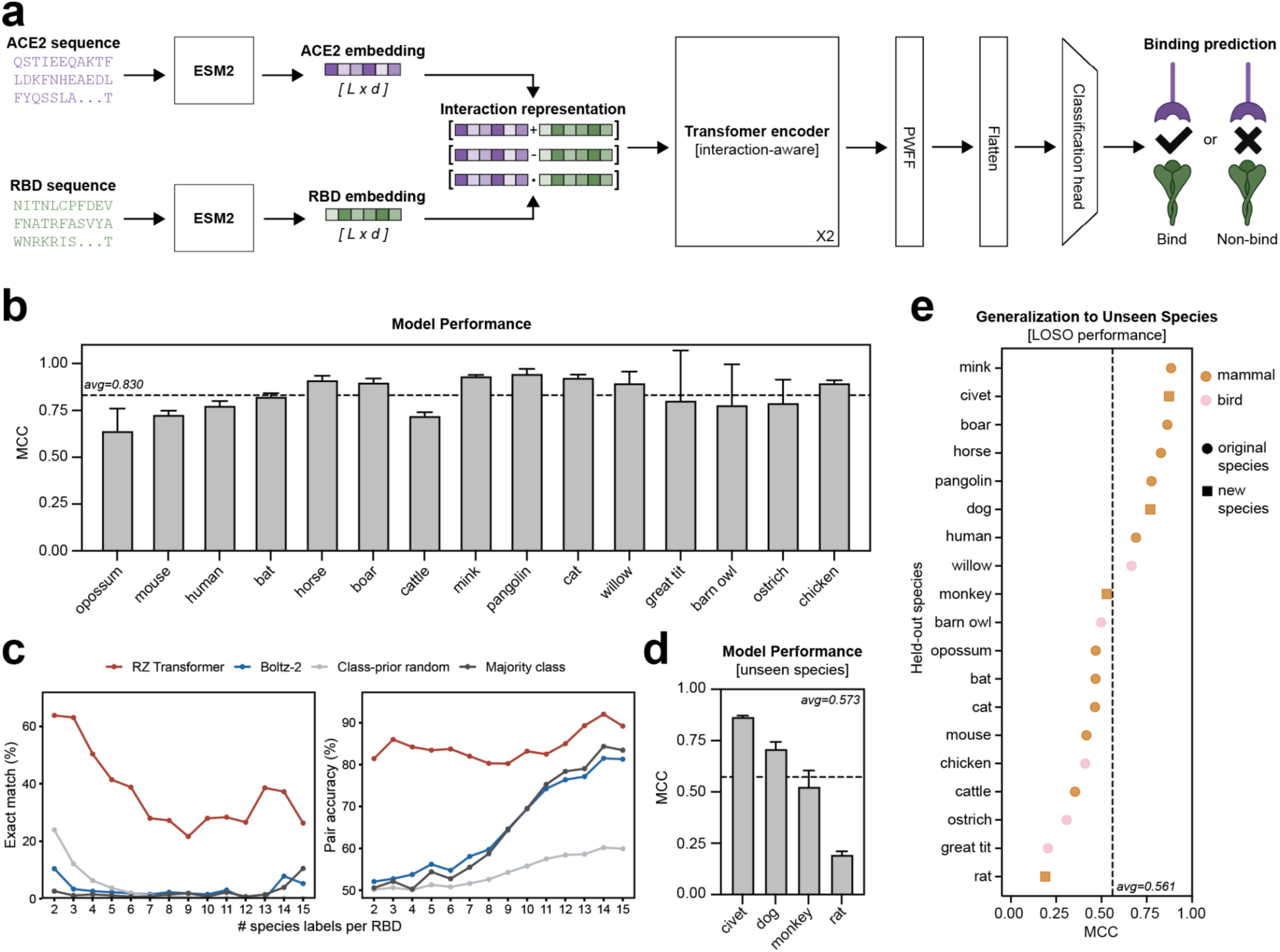
Deep learning models accurately predict RBD:ACE2 binding across diverse species. **a,** Schematic of the model architecture. ACE2 and RBD sequences are embedded using ESM2 and combined into an interaction-aware representation, which is processed by a transformer encoder followed by a classification head to predict binding. Created with <u>BioRender.com</u>. **b,** Model performance for each animal ACE2 ortholog, measured by Matthew’s correlation coefficient (MCC). Black dashed line indicates average performance across all species. **c,** Exact match rate (left) and per-interaction accuracy (right) as a function of the number of ACE2 labels per RBD variant, for the transformer, Boltz-2, and majority-class and class-prior random baselines. Exact match is the proportion of RBD variants for which all available labels were predicted correctly. Per-interaction accuracy is the proportion of individual RBD–ACE2 labels predicted correctly. Boltz-2 does not output a binding probability for protein–protein complexes, so its calls were derived from the interface confidence of the co-folded RBD:ACE2 structure, using an interface ipTM threshold of 0.76 calibrated on held-out training pairs. Interface ipTM separated experimental binders from non-binders only weakly (AUC 0.601), which accounts for its performance here. **d,** Zero-shot model performance on animal ACE2 orthologs not included in the initial screen, measured by MCC. Black dashed line indicates average performance across all species. **e,** Leave-one-species-out (LOSO) evaluation showing zero-shot model generalization to unseen animal species. For each species, all data from that species are excluded during training and used only for evaluation. Y-axis shows the test set MCC, and the x-axis indicates the species held out during training.

To further evaluate model performance, we assessed the exact match rate – the proportion of RBD sequences for which all experimentally determined ACE2 binding labels were predicted correctly. This stringent metric requires the model to correctly reconstruct the complete available binding profile for a given RBD variant and is particularly demanding for variants with mixed binding profiles, which require resolution of species-specific differences in ACE2 engagement. Exact match also becomes increasingly difficult as the number of evaluated species increases, since a single incorrect prediction causes the entire profile to be scored as incorrect.

To contextualize performance under this stringent metric, we compared the transformer with naive baselines as well as Boltz-2, a state-of-the-art structure-based co-folding model (Passaro et al., 2025). Boltz-2 provides a meaningful benchmark as it represents a general-purpose approach for evaluating protein-protein interactions directly from the sequences of an RBD:ACE2 pair, without task-specific training on our experimental binding data. For each RBD:ACE2 pair, Boltz-2 was used to co-fold the two proteins, and interface predicted TM-score (ipTM) was used as the binding proxy, with an ipTM threshold of 0.76 calibrated on held-out training pairs. Across the held-out test set, the transformer achieved an exact match rate of 94.3% (**Extended Data Fig. 3c**), and consistently outperformed Boltz-2 and the naive baselines across the full range of label counts (**Fig. 4c, left**). Although the exact match rate declined as more ACE2 labels were considered, the transformer still correctly predicted the complete 15-species binding profile for 26.6% of RBD variants, compared to 0.07% and 5.1% for random sampling and Boltz-2 respectively (**Fig. 4c, left**).

We next asked whether the decline in exact match rate reflected reduced accuracy of the individual binding predictions as label count increased. Per-interaction accuracy – the proportion of individual RBD:ACE2 calls predicted correctly – was 83.3% overall and remained above 80% across all label counts (**Fig. 4c, right**), indicating that individual binding predictions remained accurate even when complete profiles were challenging to reconstruct. By contrast, Boltz-2 achieved an overall per-interaction accuracy of 65.2%, with performance that closely tracked the majority-class baseline across label counts (**Fig. 4c, right**), suggesting that much of its accuracy reflected prediction of the dominant binding class rather than resolution of species-specific binding differences. Together, these results show that our sequence-based model trained on large-scale experimental interaction data accurately predicts RBD:ACE2 compatibility across species, consistently outperforming naïve baselines and a state-of-the-art structure-based approach.

### RBD:ACE2 binding model generalizes to unseen ACE2 orthologs

Having demonstrated that our transformer-based model could accurately predict ACE2 binding for unseen RBD variants across the 15 experimentally screened species, we next asked whether this framework could be extended to ACE2 orthologs not included in the original screen. Because ACE2 orthologs share evolutionary and sequence relationships, we reasoned that the model might transfer interaction rules learned from the original panel to predict RBD:ACE2 interactions for newly introduced receptors in a zero-shot setting.

To test this, we experimentally screened the same Omicron BA.1 RBD library against four additional ACE2 orthologs from monkey, civet, dog, and rat (**Extended Data Fig. 4a,b**) – none of which had been included during model training. We then used the resulting binding and non-binding labels as an external test set for our pretrained transformer model, without retraining or fine-tuning. Despite the challenging nature of this out-of-distribution setting, the model achieved a mean MCC of 0.573 (range 0.865 - 0.194) across the four orthologs (**Fig. 4d**), demonstrating meaningful zero-shot transfer to unseen host receptors.

To more comprehensively evaluate the generalization capacity of our model, we next performed a leave-one-species-out (LOSO) analysis. In this evaluation procedure, each species is held out for testing individually, while the model is trained on the remaining 18, cycling systematically through the dataset. LOSO represents the most stringent test of generalization available for this dataset, as it ensures that every species is treated as completely unseen during training, allowing a detailed assessment of zero-shot performance across all 19 species.

LOSO analysis demonstrated that interaction rules learned across species meaningfully transferred to ACE2 orthologs absent from training, achieving an average MCC of 0.561 across all 19 orthologs (**Fig. 4e**). Generalization was particularly strong for species including mink (0.884), civet (0.873), and boar (0.863), where the model achieved near in-distribution performance despite the receptor being entirely unseen (**Fig. 4e**). Performance was weaker for others, including great tit (0.204) and rat (0.189) (**Fig. 4e**), indicating that the capacity to generalize depends on properties of the held-out receptor. Taken together, these results establish that a model trained on a limited panel of experimentally measured ACE2 orthologs can predict RBD:ACE2 binding for receptors it has never encountered – in some cases even approaching the performance achieved when those receptors were included in training.

### Uncertainty-aware modeling prioritizes candidate ACE2-compatible species

Hundreds of species have ACE2 sequence information available, yet experimental RBD binding data exist for only a small subset, limiting assessment of their potential as candidate hosts. Although our deep learning framework can, in principle, predict interactions for any RBD:ACE2 pair, zero-shot performance varied substantially across species (**Fig. 4e**), highlighting the need for a framework that not only generates predictions, but also estimates their reliability *a priori*.

Machine learning models often degrade unpredictably when applied beyond their training distribution (Koh et al., 2020; Ovadia et al., 2019). Thus, we sought sequence-derived features that could indicate how reliably the model would generalize to an ACE2 ortholog from a new species, using leave-one-species-out (LOSO) performance across the 19 screened orthologs as an empirical benchmark. We reasoned that predictions would be more reliable for sequence pairs resembling those represented during training, than for pairs lying in more distant regions of sequence space. To test this, we concatenated the RBD and ACE2 sequence embeddings for each interaction and calculated the average distance between each held-out species and the corresponding training distribution in this joint embedding space. Average interaction embedding distance exhibited a significant negative correlation with LOSO performance (r = −0.537; p = 0.018) (**Fig. 5a**). A receptor-level measure based on mean ACE2 amino acid sequence identity between each held-out species and the training orthologs yielded a similar association (r = 0.51; p = 0.025) (**Extended Data Fig. 4c**). Hence, by both measures, the model generalized more reliably to sequences that more closely resembled the training panel.

**Fig. 5.**
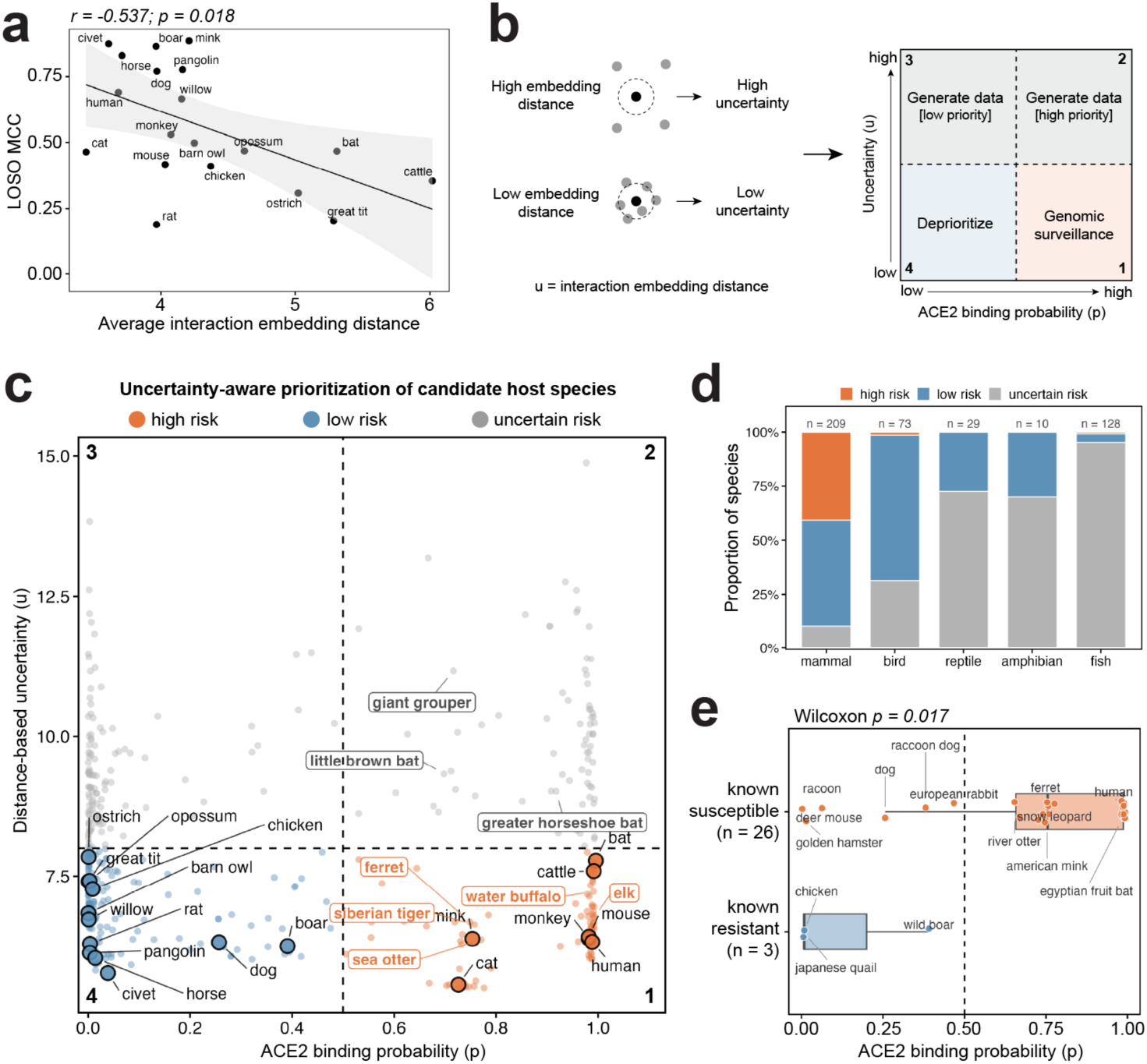
Uncertainty-aware modeling prioritizes candidate host species. **a,** Relationship between LOSO model performance (quantified by MCC), and average interaction embedding distance for each ACE2 ortholog. Spearman correlation coefficient and p-value are shown. **b,** Schematic of the uncertainty-aware prioritization framework, which combines predicted ACE2 binding probability and distance-based uncertainty to stratify species into four categories for a given SARS-CoV-2 variant: Group 1, high predicted binding and low uncertainty, prioritized for genomic surveillance; Group 2, high predicted binding and high uncertainty, prioritized for additional experimental or training data; Group 3, low predicted binding and high uncertainty, indicating under-sampled interaction space where additional data may improve model coverage; and Group 4, low predicted binding and low uncertainty, deprioritized as unlikely ACE2-compatible hosts. **c,** Predicted ACE2 binding probability for the SARS-CoV-2 Omicron BA.1 RBD (x-axis), plotted against distance-based uncertainty (y-axis) for 449 candidate host species spanning all vertebrate classes. Three of original 452 sequences were excluded as embedding distance outliers. The vertical dashed line indicates the model probability threshold for predicted ACE2 binding (p = 0.5), and the horizontal dashed line the uncertainty threshold used to stratify low- and high-confidence predictions (u = 8). Large outlined points, the 19 experimentally screened species; boxed labels, selected candidates. Species are colored according to the prioritization framework defined in b. Orange, high risk; blue, low risk; grey, uncertain risk. **d,** Risk category by vertebrate class. **e,** Predicted binding probability for species with documented SARS-CoV-2 susceptibility or resistance, restricted to confident predictions (u ≤ 8). Box, median and interquartile range; points, individual species.

To translate this observation into an actionable framework for animal prioritization, we combined joint embedding distance (as a proxy for model uncertainty) with predicted RBD binding probability (**Fig. 5b**). This yields a two-dimensional map stratifying species into four categories with distinct surveillance and data-generation priorities. Species with high predicted binding probability and low distance-based uncertainty represent the highest priority candidates for genomic surveillance (Group 1), whereas high probability predictions made with high uncertainty identify species where additional experimental or training data are most urgent (Group 2). Low probability species with high uncertainty identify lower priority data gaps that could nonetheless broaden model coverage (Group 3), while low probability species with low distance-based uncertainty can be deprioritized as unlikely ACE2-compatible hosts for that viral variant (Group 4) (**Fig. 5b**).

We applied this framework to the Omicron BA.1 RBD across 449 vertebrate ACE2 sequences, most of which lacked experimental ACE2 binding measurements. For each species, we predicted binding probability for the Omicron BA.1 RBD and calculated its distance from the interaction embedding space represented during training (**Fig. 5c**). To first assess whether the uncertainty axis behaved as intended, we examined how risk assignments were distributed across vertebrate classes. Mammals, which were well represented in the training panel, were rarely classified as uncertain (10.0%) (**Fig. 5d**). Conversely, 70.0% of amphibians, 72.4% of reptiles, and 95.3% of fish were classified as uncertain – none of which were represented during training. Birds occupied an intermediate position, with 31.5% classified as uncertain, consistent with their low representation in the training panel (5 of 19 orthologs) (**Fig. 5d**).

Among the 19 experimentally screened species, predicted Omicron BA.1 binding correctly separated binders from non-binders (**Fig. 5c**). Beyond this panel, several high-probability, low-uncertainty (Group 1) predictions corresponded to species with independent evidence of SARS-CoV-2 susceptibility, including ferret (Kim et al., 2020; Schlottau et al., 2020; Shi et al., 2020), Siberian tiger (Heniff et al., 2024; McAloose et al., 2020; Wang et al., 2026) and elk (Boggiatto et al., 2024) (**Fig. 5c**). More systematically, among species receiving confident predictions, those with published evidence of SARS-CoV-2 susceptibility had significantly higher predicted binding probabilities than the limited number of species documented as resistant (**Fig. 5e**). The model also categorized other, poorly surveyed animal species as having high binding probabilities and low uncertainty such as sea otter and water buffalo (**Supplementary Table 2**), highlighting the ability of the approach to identify unexpected candidate hosts that may otherwise be overlooked.

Group 2 contained 78 species with high predicted binding probability but high uncertainty, most of which were fish (n = 64). Of the 10 mammals in this group, 9 were bats, including several *Rhinolophus* species (**Fig. 5c**), a genus that includes important natural reservoirs of sarbecoviruses. Thus, beyond extending predictions to additional species, the uncertainty-aware framework highlighted both unsampled vertebrate classes and broader bat diversity as priorities for experimental follow-up. Together, these results establish a deep learning framework for mapping variant-specific RBD compatibility across species, translating sequence-based predictions into actionable guidance for surveillance and experimental prioritization.

## DISCUSSION

Here we present a generalizable platform that integrates high-throughput experimental screening with deep learning to predict viral host range directly from sequence. By screening a yeast display combinatorial SARS-CoV-2 RBD library across a diverse panel of animal ACE2 proteins, we generated a high-resolution map of cross-species receptor engagement and trained deep learning models that not only accurately predict RBD:ACE2 binding across the screened species, but also generalize to unseen SARS-CoV-2 variants and to animal species that were never experimentally tested. Because predictions require only an RBD and ACE2 sequence, the platform enables near-instantaneous estimation of spillover risk for emerging viral variants across the animal kingdom, serving as an “early warning system” that complements traditional genomic surveillance and field-based monitoring.

Traditional approaches to define viral host range rely on two main strategies. Experimental studies typically rely on brute-force testing of viral variants across a small panel of animal species (Li et al., 2024; Munster et al., 2020; Santini and Edwards, 2020; Shi et al., 2020; Sia et al., 2020). While informative, this approach: (i) does not scale to the vast diversity of potential hosts, (ii) cannot keep pace with the rapid emergence of new variants, and (iii) is fundamentally retrospective – variants are tested only after they are already circulating. By contrast, computational approaches can make predictions for many species in parallel and have provided valuable prospective insights (Babayan et al., 2018; Damas et al., 2020; Guo et al., 2021; Mock et al., 2021). Yet these models often lack an experimental basis, and even the most advanced computational methods treat infectivity as a static property of a virus. Importantly, with SARS-CoV-2 and other viral pathogens, host susceptibility can shift dramatically between variants (Dinnon et al., 2020; Halfmann et al., 2022). For instance, mice are resistant to early SARS-CoV-2 strains such as Wuhan-1, but are readily infected by the Omicron-lineage variants (Dinnon et al., 2020; Halfmann et al., 2022). Such observations highlight that cross-species infectivity is not fixed but can be gained or lost, demanding models with both species-level and variant-level resolution. Our platform addresses this need by training AI models on high-throughput experimental interaction data that span a broad RBD:ACE2 landscape, enabling an instantaneous spillover risk assessment for any RBD:ACE2 pair. This unified framework delivers predictions with a level of granularity not achievable by existing methods.

In prior work, we linked screening of SARS-CoV-2 RBD mutagenesis libraries with deep sequencing and machine learning to predict how combinatorial mutations impact human ACE2 binding and antibody escape – an approach we refer to as deep mutational learning (DML) (Frei et al., 2025; Shlesinger et al., 2025; Taft et al., 2022). DML has been a powerful framework for advancing prospective immunology, but current implementations treat the screening context as static. The protein used for selection (e.g., human ACE2, a monoclonal antibody, or serum) simply represents a classification label, held constant while the model learns the mapping from viral mutations to binding outcomes (Frei et al., 2025; Shlesinger et al., 2025; Taft et al., 2022). While effective, this strategy ignores a wealth of useful biological information encoded in the sequence of the interaction partner itself, effectively discarding half of the molecular recognition problem and limiting predictions to screening partners used experimentally. Recent work across molecular recognition problems has shown that jointly representing both interaction partners can support accurate prediction even when experimental data are limited (Lee et al., 2026; Reddy, 2026). Likewise, here we embed both RBD and ACE2 sequences with ESM2 and use these representations to train an interaction-aware model able to predict binding. This architecture enables predictions for any RBD:ACE2 sequence pair, including from ACE2 orthologs from species not experimentally screened.

Still, most learning algorithms are prone to failure when extrapolating beyond their training distribution. Hence, for AI models to be practically deployed, accuracy alone is not sufficient – they must also communicate if a specific output can be trusted. To address this, we integrated a distance-based uncertainty metric into our framework to quantify gaps in model knowledge during prediction. Critically, this allows us to convert raw RBD:ACE2 prediction outputs into a practical decision-making framework with four actionable categories that can directly guide genomic surveillance and experimental priorities. By distinguishing between strong, reliable predictions and speculative ones, uncertainty estimation transforms the platform into a powerful public health tool, enabling hundreds of animals to be triaged by predicted risk, guiding where time-intensive genomic surveillance and new experimental data would be most impactful.

This study has several limitations. Most notably, host-receptor engagement is a primary – but not the exclusive – determinant of viral host range. Productive infection additionally depends on entry co-factors and protease usage, tissue- and cell-type–specific receptor expression, intracellular replication competence, within-host immunity, and ecological factors that influence transmission (V’kovski et al., 2021). Therefore, a high-affinity RBD:ACE2 binding interaction indicates elevated potential for spillover, but does not guarantee infectivity or subsequent animal-to-animal transmission. Future work should integrate orthogonal data streams (e.g., host protease profiles, ACE2 expression atlases, single-cell permissivity, viral fitness and immune-escape measurements) to enable models to account for this additional biological complexity. Furthermore, our experimental ACE2 panel only contained mammals and birds, reflecting their known susceptibility to coronavirus infection (V’kovski et al., 2021). Other vertebrate classes were not represented. Similarly, the mutagenesis library was constructed in an Omicron BA.1 background (Frei et al., 2025), and as SARS-CoV-2 continues to evolve, additional libraries in more contemporaneous genetic backgrounds will be needed to keep pace with viral evolution. Nevertheless, the framework is inherently dynamic. By quantifying uncertainty, the model naturally highlights under-sampled regions of RBD:ACE2 interaction space, directing data collection on both the viral and host side to maximize information gain. As these data are incorporated, the model can be continually updated, creating an active learning cycle that steadily expands coverage and improves alongside viral evolution.

Ultimately, this AI-guided framework offers a practical and generalizable early warning system for animal reservoirs that could drive future waves of viral infection. It can forecast viral host range directly from sequence, prioritize scarce genomic surveillance resources, and improve dynamically as new data are incorporated. Though introduced here with SARS-CoV-2, the approach is broadly adaptable to other viral families, providing a blueprint for an AI-guided, One Health strategy to anticipate – and potentially prevent – future epidemics.

## MATERIALS AND METHODS

### ACE2 sequence collection, curation, and visualization

A panel of ACE2 protein sequences was assembled by combining orthologs from the NCBI and UniProt databases. For NCBI, sequences were retrieved from the “Gene” database using the query “ACE2 angiotensin converting enzyme 2” on August 8, 2022. Entries with missing amino acid sequences or substantial truncations were excluded. For UniProt, protein sequences were obtained via a search for “ace2” on August 31, 2022. Sequences flagged as “low quality,” inferred solely from homology, or containing extended stretches of unknown amino acids were removed during manual curation. The final dataset consisted of 452 unique, high-quality ACE2 protein sequences (**Supplementary Table 1**). Sequences were concatenated into a single FASTA file and aligned using MAFFT (v7.490) within Geneious (v2023.0.4), with gap padding applied to ensure uniform sequence length. The aligned sequences were one-hot encoded and flattened using a custom Python (v3.9.19) script. Principal component analysis (PCA) was performed on the resulting feature matrix, and the first two principal components were visualized using a custom R (v4.2.1) script with ggplot2 (v3.5.2).

### Phylogenetic analysis of ACE2 sequences

A maximum likelihood phylogenetic tree was generated using IQ-TREE (v2.4.0). The input was a MAFFT-aligned FASTA file containing ACE2 protein sequences from mammals and birds. IQ-TREE was run with default parameters to generate a treefile in Newick format. The resulting tree was visualized in R (v4.2.1) using the ggtree (v3.6.2) package.

### ACE2 proteins for yeast display screening

Recombinant ACE2 proteins for yeast display binding assays were purchased from commercial sources. Human (AC2-H82E6), crab-eating macaque (AC2-C52H7), rat (AC2-R5246), and masked palm civet (AC2-P5248) ACE2s were purchased from ACROBiosystems. Mouse ACE2 was purchased from Sino Biological (50249-M03H). The remaining ACE2 proteins were custom-ordered from Sino Biological. Specifically, recombinant extracellular domains were expressed in HEK293 cells by transient transfection and purified via protein A affinity chromatography. Each construct was cloned with a C-terminal human IgG1 Fc tag to promote dimerization, facilitate purification, and enable detection. Constructs included grey short-tailed opossum (*Monodelphis domestica*, NCBI protein accession: XP_007500935.1, residues 18-737), intermediate horseshoe bat (*Rhinolophus affinis*, NCBI protein accession: QMQ39227, residues 18-738), horse (*Equus caballus*, NCBI protein accession: XP_001490241.1, residues 18-738), wild boar (*Sus scrofa*, NCBI protein accession: XP_020935033.1, residues 18-739), cattle (*Bos taurus*, NCBI protein accession: XP_005228485.1, residues 18-737), american mink (*Neogale vison*, NCBI protein accession: XP_044091952.1, residues 18-739), cat (*Felis catus*, NCBI protein accession: XP_044906239.1, residues 18-739), sunda pangolin (*Manis javanica*, NCBI protein accession: XP_017505752.1, residues 20-739), willow flycatcher (*Empidonax traillii*, NCBI protein accession: XP_027757151.1, residues 18-740), great tit (*Parus major*, NCBI protein accession: XP_015486815.1, residues 23-745), barn owl (*Tyto alba*, NCBI protein accession: XP_042644640.1, residues 18-740), southern ostrich (*Struthio camelus australis*, NCBI protein accession: XP_009667495.1, residues 18-738), and chicken (*Gallus gallus*, NCBI protein accession: XP_416822.3, residues 18-740. Protein identity and integrity were verified by Coomassie-stained SDS-PAGE under reducing and non-reducing conditions (**Extended Data Fig. 1**).

### Construction of a SARS-CoV-2 Omicron RBD library for yeast display

A previously described SARS-CoV-2 Omicron BA.1 receptor-binding domain (RBD) library was used, which was constructed for yeast surface display using synthetic ssODNs with single and double NNK codon mutagenesis across tiled fragments of the RBD. Library assembly, cloning into the pYD1 vector, and transformation into *Saccharomyces cerevisiae* strain EBY100 were performed exactly as previously reported(Frei et al., 2025).

### Screening RBD libraries for ACE2 binding

Yeast cells containing the Omicron RBD library plasmid were cultured in SD-CAA medium for 18–24 hours at 30°C with mild agitation. To induce RBD surface display, cells were transferred into SG-CAA medium [20 g/L galactose (Sigma, G0625), 8.56 g/L NaH₂PO₄·H₂O (Roth, K300.1), 6.77 g/L Na₂HPO₄·2H₂O (Sigma, 1.06580), 6.7 g/L yeast nitrogen base without amino acids (Sigma, Y0626), and 5 g/L casamino acids (Gibco, 223120) in deionized water] and incubated at 23°C for 24 hours. Following induction, 40 million yeast cells were harvested by centrifugation at 8000 RPM for 3 minutes and washed twice in Wash Buffer [0.5% bovine serum albumin (Sigma, A2153), 2 mM EDTA (Biosolve, 051423), 0.1% Tween-20 (Sigma, P9416) in PBS (Pan Biotech, P04-53500)]. A concentration of 50nM ACE2 has been established in prior studies as sufficient to detect physiologically relevant binding interactions (Frei et al., 2025; Taft et al., 2022). Reported affinities between ACE2 and natural SARS-CoV-2 RBD variants typically fall within the single- to double-digit nanomolar range. For instance, Wuhan (27.5nM), Alpha (6.7nM), Beta (19.7nM), Gamma (16.0nM), Delta (25.1nM), and Omicron (5.5nM)(Kim et al., 2023; Li et al., 2022). Therefore, RBD variants detectable at 50nM ACE2 are expected to exhibit binding affinities consistent with those of infectious SARS-CoV-2 strains and within a physiologically meaningful range. Thus, ACE2 proteins were used at a final concentration of 50nM for mouse, human, intermediate horseshoe bat, cattle, and crab-eating macaque. For ACE2 orthologs that showed minimal binding to the naïve library under these conditions (grey short-tailed possum, horse, wild boar, american mink, cat, sunda pangolin, willow flycatcher, great tit, barn owl, southern ostrich, chicken, dog, rat, masked palm civet), the concentration was increased to 500nM to improve sensitivity and capture lower-affinity or rare binders that may still represent biologically relevant interactions. Thus, binder proportions are not directly comparable across species. Cells were stained with the ACE2 orthologs rotating overnight at 4°C. Following a single wash, cells were treated with 1:100 diluted anti-human IgG AF647 (Jackson ImmunoResearch, 109-605-098) or 1:40 diluted anti-His AF647 (Qiagen, 35370) to detect ACE2 binding, and incubated rotating for 1 hour at 4°C. After two washes, cells were stained with 1:150 diluted anti-FLAG PE (BioLegend, 637310) to detect surface expression, and incubated rotating for 30min at 4°C. Subsequently, cells were washed and resuspended in Wash Buffer for fluorescence-activated cell sorting (FACS) on a BD Influx, BD FACSAria Fusion, or BD FACSDiscover S8. Cells were sorted into ACE2-binding (PE⁺/AF647⁺) and non-binding (PE⁺/AF647⁻) populations. Sorted populations were pelleted at 3000 RPM for 10 minutes and resuspended in SD-CAA medium for outgrowth at 30°C. Multiple sorts were performed per species to enrich each population.

### Deep sequencing of RBD libraries

The pYD1 plasmid encoding the RBD library was extracted from yeast cells according to the instructions of the manufacturer (Zymo, D2004). Subsequently, two consecutive PCR reactions were performed to prepare the library for deep sequencing. The first PCR mixture contained NEBNext Ultra II Q5 Master Mix (New England Biolabs, M0544X), 80ng of template DNA, and custom designed primers which possessed a 0-3bp heterogeneity spacer (Fadrosh et al., 2014), and Illumina partial adaptor sequences. PCR 1 conditions were: initial denaturation at 98°C for 5 minutes, followed by 15 cycles of denaturation at 98°C for 30 seconds, annealing at 64°C for 90 seconds, extension at 72°C for 35 seconds, and a final extension at 72°C for 6 minutes. PCR products were cleaned-up using SPRIselect beads (Beckman Coulter, B23318) according to the instructions of the manufacturer. For the second PCR, the mixture contained NEBNext Ultra II Q5 Master Mix (New England Biolabs, M0544X), 20ng of template DNA, and Nextera forward and reverse indexing primers (Illumina, 20027215). PCR 2 conditions were: initial denaturation at 95°C for 3 minutes, followed by 20 cycles of denaturation at 98°C for 20 seconds, annealing at 62°C for 25 seconds, extension at 72°C for 35 seconds, and a final extension at 72°C for 6 minutes. PCR products were cleaned-up using SPRIselect beads (Beckman Coulter, B23318). Purified amplicons were pooled and assessed for sample quality using a Bioanalyzer, then subjected to paired-end sequencing using a MiSeq v3 Reagent Kit (600 cycles) (Illumina, MS-102-3003) or an Aviti High Output Kit (600 cycles) (Element Biosciences, 860-00008).

### Preprocessing of deep sequencing data

Paired-end sequencing reads were paired, trimmed, and merged using the BBTools suite, with a Phred quality threshold of Q > 20. RBD nucleotide sequences were then extracted using custom Python (v3.9.19) scripts, and additional downstream preprocessing was performed in R (v4.2.1). Extracted nucleotide sequences were translated, and sequences containing non-ATCG characters, frameshifts, or premature stop codons were removed. Wild-type Omicron BA.1 RBD sequences and variants supported by fewer than two sequencing reads were also excluded. Duplicate amino acid sequences were collapsed within each sorted population, and variants detected in both ACE2-binding and non-binding populations for the same ACE2 ortholog were removed to avoid conflicting labels. Remaining variants were assigned binary binding labels according to their sorted population.

### Interaction dataset and feature engineering

RBD and ACE2 amino acid sequences were embedded using ESM2 (8M) (Lin et al., 2023). The use of a pre-trained protein language model enabled extraction of functionally informative sequence representations and projection of RBD and ACE2 sequences into a shared latent space prior to supervised learning. For each RBD pair, RBD embeddings were retained at residue-level resolution, whereas ACE2 residue embeddings were mean-pooled across non-special tokens to generate a receptor-level representation. Because the number of ACE2 species was limited (∼15), learning ACE2-specific representations ab initio would likely be data limited. We therefore used the pooled ACE2 embedding as a biologically informed contextual representation, expanding it across the RBD sequence length. At each RBD position, a pairwise feature vector was constructed by concatenating: (i) the RBD residue embedding, (ii) the absolute difference between the RBD residue embedding and the pooled ACE2 embedding, and (iii) the element-wise product between the RBD residue embedding and the pooled ACE2 embedding. These operations were designed to emphasize subtle interaction-level differences between sequence representations, which may not be sufficiently resolved from the original ESM embeddings alone.

### RBD:ACE2 transformer model architecture

Interaction features were passed through a linear projection layer, followed by learned positional embeddings. The projected features were then passed to a transformer encoder with padding masks applied to padded RBD positions. The encoder output was passed through a position-wise feedforward network that learned how to aggregate the residue-level embeddings into a compact representation by projecting them to a lower-dimensional space. The resulting representations were subsequently flattened across sequence positions and passed to a final linear classification head. The model output was a single logit for each RBD:ACE2 pair. Predicted binding probabilities were obtained by applying a sigmoid transformation to the logit. The model was trained using binary cross-entropy loss.

### Model training and evaluation

RBD variants were first clustered to reduce sequence-level leakage between training and test sets. Amino acid sequences were one-hot encoded and clustered using k-means clustering, yielding five RBD sequence clusters. Two clusters were assigned to the training set, while the remaining clusters were reserved for validation and testing, corresponding to a 90.9% training split and a 9.1% test split. Clusters used for training and testing were selected to maintain a balanced train/test ratio and sufficient numbers of binders and non-binders for each species across partitions This cluster-based splitting strategy was used to limit overlap of closely related RBD variants across partitions and to evaluate generalization to RBD variants with distinct mutational patterns. Model hyperparameters were selected by iterative grid search over architecture and training parameters, including latent embedding dimension, number of transformer encoder layers, number of attention heads, transformer feedforward dimension, position-wise feedforward dimension, dropout rate, learning rate, batch size, and number of training epochs. Finally, the model trained with learning rate 0.0001, for 5 epochs using a batch size of 32. No gradient clipping was applied. Candidate models were trained on the training set and selected based on validation-set MCC performance with early stopping after 2 epochs, and final performance was reported on the held-out test set. Dropout was applied after positional encoding, within the transformer encoder layers, and after the position-wise feedforward module during training to prevent overfitting. ESM2 was used to generate sequence representations for RBD and ACE2 sequences, with ESM2 parameters kept frozen during training. Model performance was assessed using Matthews correlation coefficient (MCC), calculated separately for each ACE2 ortholog and averaged across experimentally screened species. We additionally calculated the exact match rate, defined as the proportion of RBD variants for which all available ACE2 binding labels were predicted correctly. Exact match rate was calculated across available labels only. For this analysis, test-set RBD variants were stratified into uniform-label variants, defined as variants that either bound all available ACE2 orthologs or failed to bind all available ACE2 orthologs, and mixed-label variants, defined as variants that bound ACE2 from at least one species but not another. Exact match rate was calculated separately for uniform-label and mixed-label variants. All experiments were conducted on a single NVIDIA GeForce RTX 4090 GPU with an average training time of ∼4 hours.

### Feature block importance for RBD:ACE2 interaction features

The transformer combines each RBD residue embedding with the pooled ACE2 embedding through three operations, the RBD embedding itself, the absolute difference between it and the pooled ACE2 embedding, and their element-wise product. Attributing importance to individual operations inside the encoder is confounded by residue-level self-attention and by the flattening step preceding the classification head, so block importance was quantified on a simpler surrogate classifier applying the same operations to mean-pooled representations, where each operation forms a single contiguous block of input features. Mean-pooled ESM2 (8M) embeddings for RBD (r) and ACE2 (a), 320 dimensions each, were concatenated into the 1,280-dimensional vector x = [r, a, |r − a|, r · a]. The pooled ACE2 embedding was retained as an explicit fourth block, although it is not a standalone block in the transformer, so that the contribution of the unmodified receptor could be separated from the two interaction operations. The surrogate was a multilayer perceptron with two hidden layers (512 and 128 units, ReLU, dropout 0.3) and a single logit output, trained for 5 epochs with Adam (learning rate 1 × 10^-3^, batch size 2,048) under binary cross-entropy reweighted by the inverse training class ratio. Features were standardized using training-set statistics. Partitions reused the k-means RBD clusters defined for the main model, clusters 0 and 2 for training, 4 for validation and 1 and 3 for testing, so that no RBD variant was shared between partitions. Pairs were subsampled under a fixed seed to 200,000, 50,000 and 50,000 respectively to keep the repeated permutation and ablation passes tractable. The decision threshold was selected by maximizing MCC on the validation partition (0.525). At this operating point the surrogate reached test MCC 0.824 (AUC 0.970), comparable in magnitude to the held-out MCC of 0.830 reported for the transformer, which supports its use as a stand-in for assessing the relative value of the feature operations. The two values are not computed identically, as the transformer value is averaged across species and the surrogate value is pooled over all test pairs. Three complementary measures were computed, all relative to the test-partition baseline. For permutation importance, the 320 dimensions of one block were jointly shuffled across pairs and the decrease in test MCC recorded over five independent permutations. For ablation importance, one block was replaced either by zeros or by its feature-wise test-partition mean. Both ablation modes are reported because zeroing a standardized block imputes the training mean, whereas the test mean removes that block’s variation without introducing a distribution shift relative to the evaluation data. As a model-free reference, the point-biserial correlation between each standardized feature dimension and the binary binding label was computed on the training partition and summarized per block. Two constraints apply to the interpretation of these measures. Because |r − a| and r · a are deterministic functions of r and a, the four blocks are mutually redundant, so single-block permutation and ablation measure the unique contribution of each block given all others and are lower bounds that must not be read additively. The analysis was also performed on the mean-pooled surrogate rather than on the transformer, so it establishes the relative value of the four feature operations but not their absolute importance within the deployed architecture.

### Benchmarking and hyperparameter tuning

To identify the optimal encoder architecture, multiple models were trained using the same cluster-based data split strategy and input features derived from the ESM embeddings. We evaluated XGBoost, a multilayer perceptron (MLP) and a fine-tuned ESM-8M encoder as alternatives to the proposed transformer. Hyperparameters for each model were optimized by grid search under the same training and evaluation framework (average species MCC). The transformer achieved the highest performance (average MCC 0.78), outperforming XGBoost (0.66), the fine-tuned ESM-8M encoder with position-wise feed-forward (PWFF) pooling (0.629) and the MLP baseline (0.374). The final transformer used an embedding dimension of 256 matching the ESM-8M output, 8 attention heads, 4 encoder layers, a hidden feed-forward dimension of 1,024 and a position-wise feed-forward dimension of 128.

### Boltz-2 co-folding baseline for RBD:ACE2 binding prediction

To benchmark against a structure-based approach, we used Boltz-2 (Passaro et al., 2025), an open-source biomolecular co-folding model in the AlphaFold3 family. Each RBD:ACE2 pair was scored by co-folding the RBD (chain A) with the full-length ACE2 ortholog of the corresponding species (chain B) as a two-chain complex and reading the interface confidence of the top-ranked predicted structure. Boltz-2 does not output a binding affinity or probability for protein– protein complexes, as its affinity head is restricted to small-molecule ligands. The binding signal used here is therefore structural confidence, meaning how confidently the model docks the two chains into a well-defined interface, used as a proxy for binding. This is the standard way co-folding models are applied to protein-protein interaction prediction and is the only binding-relevant signal Boltz-2 provides for this class of complex. Three interface confidence metrics were extracted from the top-ranked sample (model_0). protein_iptm is the interface predicted TM-score over protein chains only (0–1, higher is better), estimating the accuracy of the predicted relative placement of the two chains, and is the primary discriminative signal. complex_ipde is the predicted distance error at the interface in Angstroms (lower is better). complex_iplddt is the predicted local distance difference test averaged over interface residues (0–1, higher is better). Values reported on a 0–100 scale were rescaled to 0–1, and missing values were treated conservatively (ipTM and ipLDDT set to 0, iPDE set to 99 Å) so that an unscored complex could never be called a binder. A pair was called a binder only when all three gates were satisfied, protein_iptm ≥ 0.76, complex_ipde ≤ 20.0 Å and complex_iplddt ≥ 0.75. The ipTM threshold was calibrated so that Boltz-2 would be evaluated at its own empirically optimal operating point rather than an arbitrary cutoff. Calibration used labelled training pairs disjoint from the evaluation set. ipTM was swept from 0.05 to 0.95 in increments of 0.01, MCC against the experimental labels was computed at each grid value, and the sweep was repeated over 10 independent random subsamples of 500 pairs under a fixed seed. The threshold maximizing the mean MCC across subsamples was selected, as averaging the MCC curves before taking the maximum is more stable than averaging the per-subsample optima. The selected value of 0.76 was also the per-subsample optimum in 8 of 10 subsamples and was applied unchanged to the held-out evaluation set. The complex_ipde and complex_iplddt cutoffs were not tuned and serve as permissive structural sanity gates intended only to reject grossly ill-defined interfaces. On this dataset both were satisfied by all folded complexes, so the binder call reduces in practice to protein_iptm ≥ 0.76 alone. Structures were predicted with diffusion_samples = 1 and recycling_steps = 3. Identical (RBD sequence, ACE2 sequence) pairs were folded once and the metrics reused. Multiple sequence alignments were computed once per unique chain sequence and referenced from every complex input, retrieved from the ColabFold MSA server (api.colabfold.com) using run_mmseqs2 and supplied as unpaired per-chain a3m files. Unpaired alignments are appropriate because cross-species RBD:ACE2 pairs share no co-evolutionary signal that chain pairing could exploit. Each alignment was keyed by the SHA-1 hash of its query sequence to guarantee correct alignment-to-sequence assignment and prevent silent fallback to a single-sequence alignment. Folding was distributed across GPUs by sharding complexes on residue count, running one boltz predict process per device. Boltz-2 was evaluated on held-out RBD variants, restricted to the 19 ACE2 orthologs with a curated full-length receptor sequence. RBD variants were sampled whole, with all labelled pairs for a given RBD kept together and stratified by the number of available ACE2 labels per variant, so that exact match rate and per-interaction accuracy could be resolved by label count exactly as for the transformer (**Fig. 4c**). Calls were scored against the same experimental labels, metrics and baselines used for the transformer.

### Leave-one-species-out (LOSO) evaluation

Leave-one-species-out evaluation was performed by iteratively withholding all RBD:ACE2 pairs corresponding to one experimentally screened ACE2 ortholog. For each iteration, model training and hyperparameter selection were performed using only pairs from the remaining ACE2 orthologs, and final evaluation was performed on the held-out species. This provided a zero-shot evaluation for each ACE2 ortholog, as no binding data from the held-out species were used during training. This procedure was repeated for each screened ACE2 ortholog. Performance was quantified using the MCC for each held-out species.

### Distance-based uncertainty estimation

Distance-based uncertainty was estimated from averaged RBD:ACE2 interaction embeddings. For each ACE2 ortholog, we paired the ACE2 sequence with each RBD sequence and generated a concatenated RBD:ACE2 embedding for every pair. These pairwise embeddings were averaged across RBD sequences to obtain a single interaction-space representation for that ACE2 ortholog. We repeated this procedure for each experimentally measured training species and then calculated Euclidean distances between each query ACE2 representation and the corresponding training-species representations. For each query ACE2 ortholog, the average Euclidean distance to the experimentally measured training-species representations was used as the distance-based uncertainty score. To characterize the relationship between interaction distance and predictive performance, distances calculated during LOSO evaluation were compared with held-out species MCC. An isotonic regression model was fit to estimate the relationship between interaction distance and LOSO performance. For downstream prioritization, an interaction-distance threshold of u = 8 was used to distinguish lower- and higher-uncertainty predictions. Predictions for additional ACE2 orthologs were then reported together with predicted binding probability and distance-based uncertainty, and stratified into lower- and higher-uncertainty groups using this threshold.

### Statistics and reproducibility

Random seeds were fixed for all classifier training procedures, so that model initialization, data partitioning and subsampling were reproducible.

## ACKNOWLEDGEMENTS

We thank Thomas Horn, Mariangela Di Tacchio, Chiara Cavallini, Svitlana Malysheva, and Aleksandra Gumienny from the Single Cell Facility at the Department of Biosystems Science and Engineering (D-BSSE), ETH Zürich, for their valuable support with flow cytometry-based screening. We are also grateful to Christian Beisel, Mirjam Feldkamp, Elodie Burcklen, and Ina Nissen at the D-BSSE Genomics Facility in Basel for their excellent assistance with next-generation sequencing.

## AUTHOR CONTRIBUTIONS

Conceptualization: E.B.I, S.T.R. Methodology: E.B.I., T.B., T.S., L.P., L.F., N.C., E.S., J.M.T. Software: T.B., T.S., L.P., B.S. Validation: E.B.I., T.B., T.S., L.P. Formal analysis: E.B.I., T.B., T.S., L.P., E.S., B.S. Investigation: E.B.I., T.B., T.S., L.P., L.F. Resources: L.F., J.H., J.M.T., S.T.R. Data curation: E.B.I., T.B., T.S., L.P., J.H. Writing – Original Draft: E.B.I., T.B. Writing – Review & Editing: all authors. Visualization: E.B.I., T.B., T.S. Supervision: E.B.I., T.B., S.T.R. Project administration: E.B.I, T.B., S.T.R. Funding: E.B.I., S.T.R.

## DATA AND MATERIALS AVAILABILITY

All data needed to evaluate the conclusions in the paper are present in the paper and/or the Supplementary Materials. The ACE2 binding dataset generated in this study has been deposited at XXX, and is publicly available. Custom code for model training and evaluation is available at: https://github.com/LSSI-ETH/reverse_zoonosis.

## EXTENDED DATA FIGURES

**Extended Data Fig. 1.**
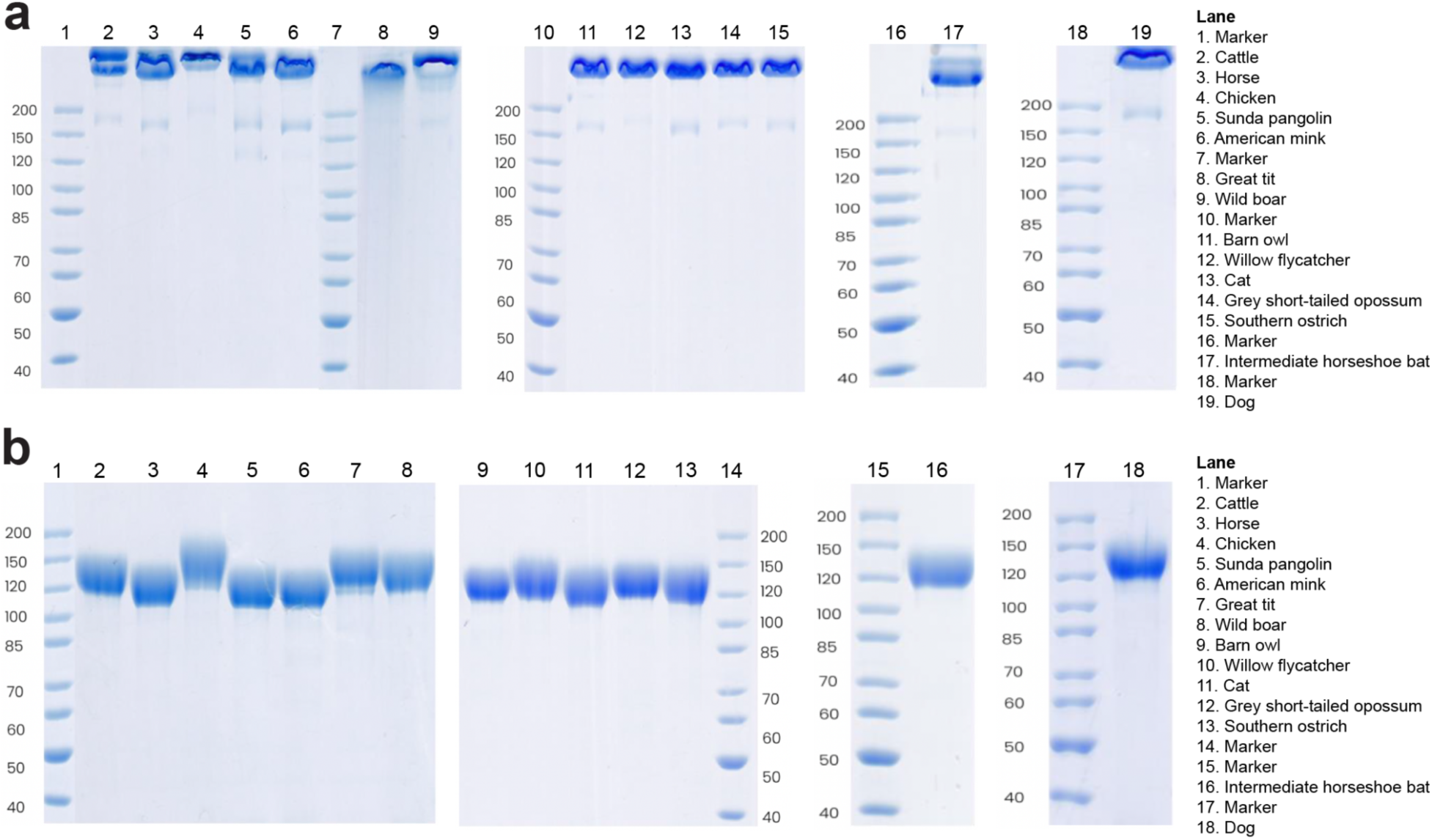
Quality control of custom-produced animal ACE2 proteins. Coomassie blue-stained SDS-PAGE gel under **a,** non-reducing conditions, and **b,** reducing conditions. Molecular weight markers are indicated in kilodaltons.

**Extended Data Fig. 2.**
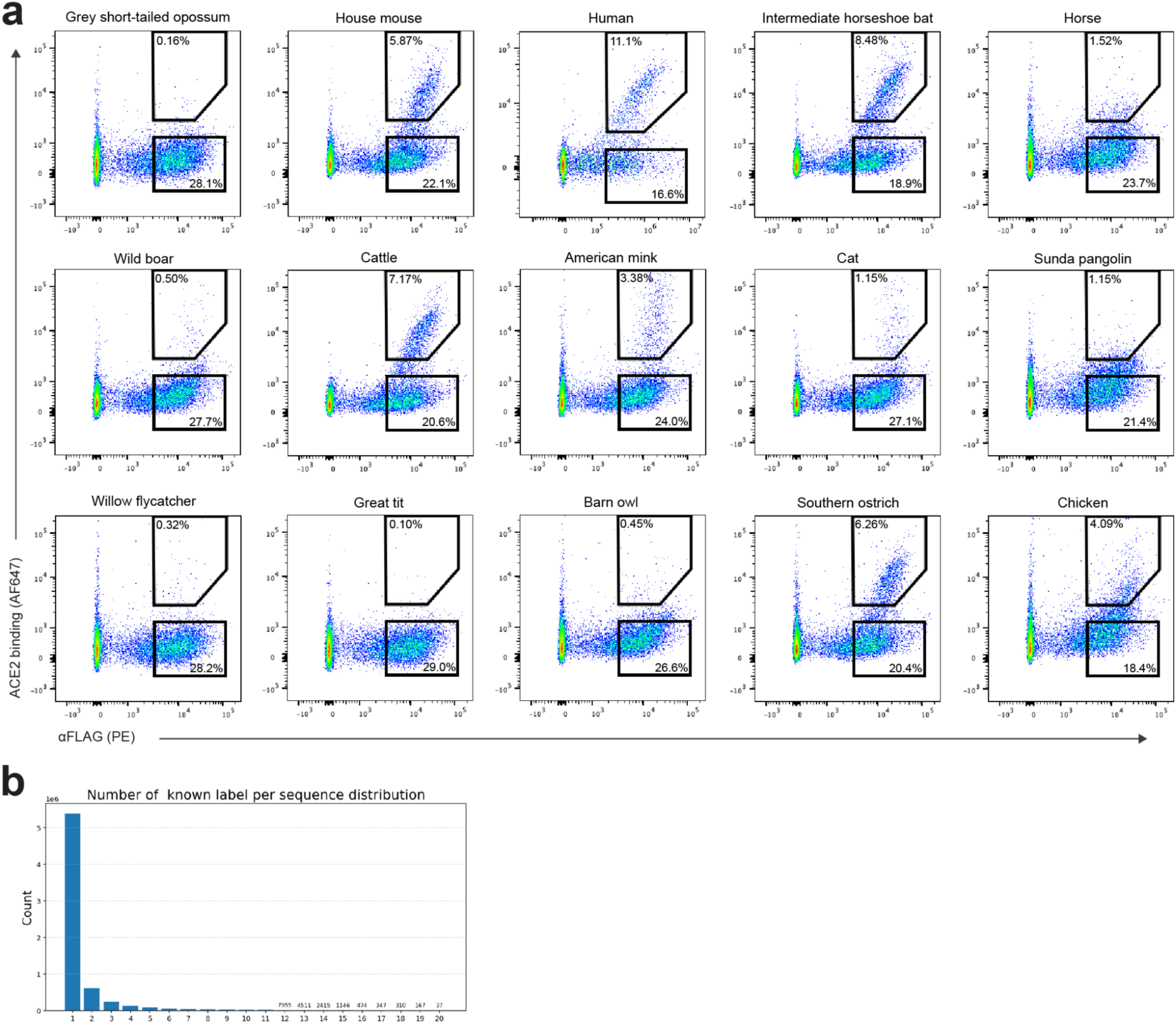
Flow cytometry analysis and sequence coverage for ACE2-binding screens. **a,** Representative flow cytometry plots of the SARS-CoV-2 Omicron BA.1 RBD yeast surface display library stained with the indicated soluble ACE2 orthologs. X-axis: surface display measured by anti-FLAG staining. Y-axis: ACE2 binding. Gated percentages indicate the proportion of ACE2-binding and non-binding populations. **b,** Distribution of the number of ACE2 binding labels assigned per RBD sequence after FACS sorting and deep sequencing.

**Extended Data Fig. 3.**
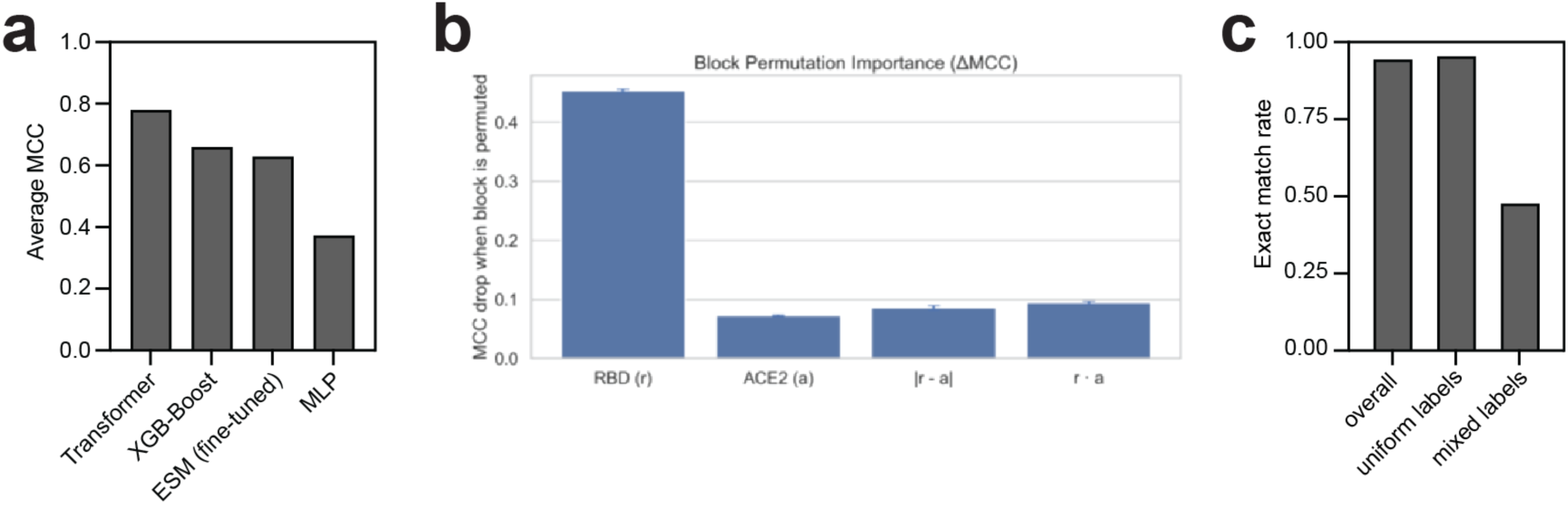
Evaluation of the RBD:ACE2 prediction framework. **a,** Benchmarking of model architectures. MCC averaged across species for each strategy. **b,** Contribution of RBD:ACE2 interaction feature blocks to binding prediction, quantified by permutation importance. Bars show the decrease in test-set MCC (ΔMCC) after jointly shuffling each feature block across pairs; error bars indicate the standard deviation across five independent permutations. **c,** Exact match rate across all ACE2 orthologs per RBD sequence – defined as the fraction of sequences for which all ACE2 binding labels were predicted correctly. Results are shown for the overall test set, as well as for RBD sequences with uniform (all the same) or mixed (at least one different) binding labels across ACE2 orthologs.

**Extended Data Fig. 4.**
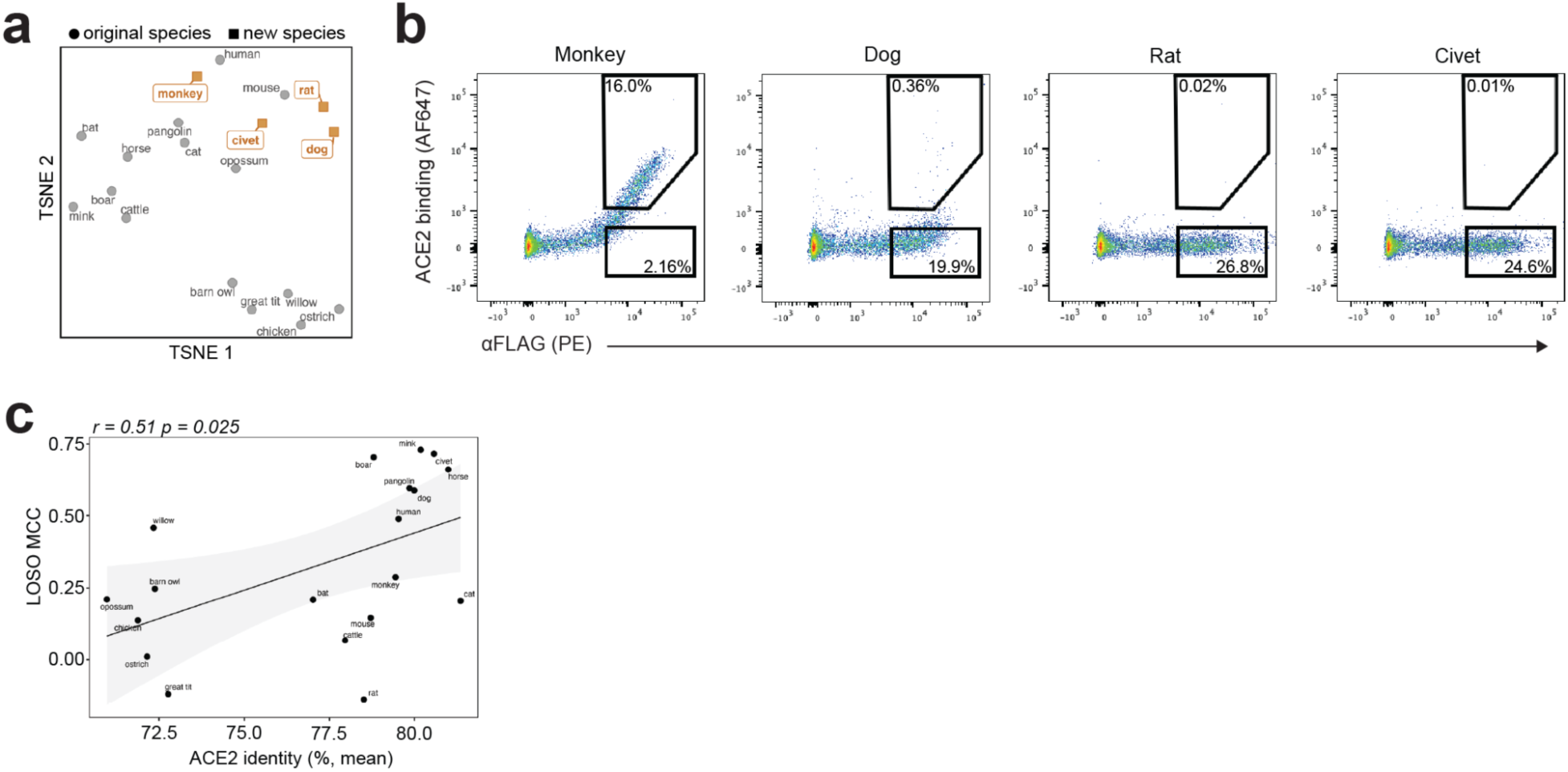
Generalization across ACE2 sequence space. **a,** TSNE visualization of ACE2 sequence embeddings generated by ESM2. Species in the original dataset are shown as grey circles, and newly added species (all mammals) as orange squares. **b,** Representative flow cytometry plots of the SARS-CoV-2 Omicron BA.1 RBD yeast surface display library stained with the indicated soluble ACE2 orthologs. X-axis: surface display measured by anti-FLAG staining. Y-axis: ACE2 binding. Gated percentages indicate the proportion of ACE2-binding and non-binding populations. **c,** Relationship between LOSO model performance and mean ACE2 amino acid sequence identity between each held-out ortholog and the training species. Pearson correlation coefficient and p-value are shown. Line, linear fit; shading, 95% confidence interval.

## SUPPLEMENTARY TABLES

**Supplementary Table 1 | Full ACE2 sequence collection.**

**Supplementary Table 2 | Prioritization of host spillover risk.**

